# Dynamic map illuminates Hippo to cMyc module crosstalk driving cardiomyocyte proliferation

**DOI:** 10.1101/2022.10.11.511763

**Authors:** Bryana N. Harris, Laura A. Woo, R. Noah Perry, Mete Civelek, Matthew J. Wolf, Jeffrey J. Saucerman

## Abstract

Cardiac diseases are characterized by the inability of adult mammalian hearts to overcome the loss of cardiomyocytes (CMs). Current knowledge in cardiac regeneration lacks a clear understanding of the molecular systems determining whether CMs will progress through the cell cycle to proliferate. Here, we developed a computational model of cardiac proliferation signaling that identifies key regulators and provides a systems-level understanding of the cardiomyocyte proliferation regulatory network. This model defines five regulatory networks (DNA replication, mitosis, cytokinesis, growth factor, hippo pathway) of cardiomyocyte proliferation, which integrates 72 nodes and 88 reactions. The model correctly predicts 72 of 76 (94.7%) independent experiments from the literature. Network analysis predicted key signaling regulators of DNA replication (e.g., AKT, CDC25A, Cyclin D/CDK4, E2F), mitosis (e.g., Cyclin B/CDK2, CDC25B/C, PLK1), and cytokinesis, whose functions varied depending on the environmental context. Regulators of DNA replication were found to be highly context-dependent, while regulators of mitosis and cytokinesis were context-independent. We also predicted that in response to the YAP-activating compound TT-10, the Hippo module crosstalks with the growth factor module via PI3K, cMyc, and FoxM1 to drive proliferation. This prediction was validated with inhibitor experiments in primary rat cardiomyocytes and further supported by re-analysis of published data on YAP-stimulated mRNA and open chromatin of Myc from mouse hearts. This study contributes a systems framework for understanding cardiomyocyte proliferation and identifies potential therapeutic regulators that induce cardiomyocyte proliferation.

## Introduction

Cardiovascular diseases are the leading causes of death worldwide and are the highest cause of morbidity and mortality (CDC, 2022). While the survival rate from cardiovascular disease has increased over the past few decades, heart failure morbidity and mortality have remained staggering. A large part of this decline is imparted by cardiomyocyte (CM) loss and no supporting CM renewal capacity. Currently, proposed therapies can only reverse or slow certain aspects of cardiac dysfunction and disease, failing to replace lost CMs. Instead, the heart undergoes wound healing to replace dead CMs with fibrotic scar tissue, which impairs normal cardiac function (Leach & Martin, 2018). Therefore, there is a strong need for therapeutic strategies that increase CM renewal and, in turn, restore the contractile function of the heart and considerably improve survival and disease outcomes (Bertero & Murry, 2018; Eschenhagen et al., 2017; Hashimoto et al., 2018; Mann & Bristow, 2005).

One promising therapeutic strategy is to induce the proliferation of endogenous cardiomyocytes (Zhao et al., 2020) by manipulating signaling molecules and regulatory proteins. This strategy will require an understanding of both the molecular mechanisms that promote cardiomyocyte cell division and those that result in a cell cycle exit (Payan et al., 2020). Over the past few years, it has been found that manipulating cell cycle regulatory and inhibitory proteins can promote cardiomyocyte proliferation. Previous studies have also shown that combining four cell cycle regulators could stimulate adult cardiomyocyte proliferation *in vivo* (Mohamed et al., 2018). In addition to these cell cycle proteins, several signaling pathways, such as YAP and neuregulin signaling pathway proteins, stimulate CM proliferation (Eschenhagen et al., 2017; Hashimoto et al., 2018; Hashmi & Ahmad, 2019; Mann & Bristow, 2005; Oyama et al., 2013; Yester & Kühn, 2017; Yutzey, 2017). microRNAs have also been implicated in stimulating CM proliferation (Eulalio et al., 2012; Ouyang & Wei, 2021; Tian et al., 2015; Torrini et al., 2019) Like other complex biological systems, innovative models and quantitative analyses are needed to unravel specific pathways and crosstalk between various regulators affecting CM proliferation.

In this study, we constructed and experimentally validated what is, to our knowledge, the first computational model of cardiomyocyte proliferation, which predict molecular drivers of cell cycle progression. The model integrates five modules that incorporate multiple regulatory pathways. With a virtual knockdown screen, we predict how the influence of particular nodes changes with different signaling contexts. Additionally, the model elucidated crosstalk between the growth factor and Hippo modules. We also identified key hubs for which the Hippo signaling pathway regulates cardiomyocyte proliferation. This model provides a systems framework with which to prioritize potential signaling drivers of therapeutic modulators of CM proliferation, in which model predictions are validated with *in vitro* experimentation and *in vivo* data.

## Materials and Methods

### Model Construction

A predictive computational model of the cell cycle-proliferation signaling network in CMs was manually constructed from published literature and resources such as KEGG, SIGNOR, and Cell Signaling(*Cell Cycle Regulation Pathways*, n.d.; *KEGG PATHWAY: Cell Cycle - Homo Sapiens (Human)*, n.d.; Licata et al., 2020). The literature search began by identifying references that indicate a role for specific proteins, whether by a diagram of the activity or the role of specific proteins in the heart. Direct molecular reactions between proteins were included if multiple sources supported the interaction. However, some interactions were not found in cardiomyocytes. All papers involving in vitro or in vivo experiments performed in rat cardiomyocytes or hiPSC-CMs were reserved for validation during the literature review. Model outputs were set as general cell cycle phases that are measurable through experiments such as DNA replication (measured by EdU assay), general cell cycle activity (fluorescent staining of Ki67), mitosis (measured by phH3 positive cells), or cytokinesis (measured by Aurora B presence at the midbody of the cell). The overall network contains 72 species and 88 reactions. The model includes 14 inputs, including receptor inputs (neuregulin 1, insulin-like growth factor (IGF), and fibroblast growth factor (FGF)) and five phenotypic outputs (DNA replication, mitosis, polyploid, cytokinesis, binucleation). This network integrates five modules that increase CM proliferation: DNA replication module, mitosis module, cytokinesis module, growth factor module, and Hippo module. Full documentation supporting model reactions is provided in **Supplementary File 1**, and all references are provided in **Supplementary File 2**.

Signaling dynamics were predicted using a previously described logic-based differential equation (LDE) approach (Ryall et al., 2012). In this method, the activation of one node in the model by another is modeled using a normalized Hill function. Logical AND or OR operations were used to represent pathway crosstalk. OR gating is used when each input to a node is sufficient but not necessary for activation, whereas AND gating is used when each input is necessary. Default reaction parameters included reaction weight (w = 1), Hill coefficient (n = 1.4) and half-maximal effective concentration (EC_50_ = 0.5), and default node parameters included initial activation (Y_init_= 0), maximal activation (Y_max_ = 1) and time constant (τ = 1). The reaction weight for model inputs (Oxygen, HIPK2, AurkABora, SMAD3, AurB, Bub1, Nrg1, FGF2, IGF1, PKA, Mst1, Rho, and PRC1) was chosen to maximize or minimize the number of nodes activated between 25% and 90%, preventing a drastic increase or decrease in node activity to obtain the most information from the sensitivity analysis. The system of ODEs was auto-generated from Supplementary File 1 using Netflux (available at https://github.com/saucermanlab/Netflux) and implemented in MATLAB.

### Model Validation

The literature used to validate the network’s input-output relationships was identified by searching for each network relationship together with the term “cardiomyocyte” or “cardiomyocyte proliferation” in the Pubmed or Google Scholar database. To create quality or reproducible models, validations used were only from studies using rats, mice, or human cardiomyocytes. All supporting studies used were independent of those used to develop the model network. Validation was performed by comparing the qualitative increase, decrease, or no change in output activity of the model simulation to the experimental results. Changes of less than 0.5% were categorized as “no change.” All model validations along with supporting literature are provided in **Supplementary Document 2**.

### Sensitivity Analysis

Functional analysis of the model was performed by simulating individual knockdowns for each of the 74 species in the network and predicting the corresponding change in the activity of every other node in the network. First, the steady-state activity of each node was computed under baseline conditions (control). We then knocked down the activity of each node one a time and subtracted the basal activity values from the values in the knockdown state to calculate “Δ Activity.” Influence is measured as the number of nodes with a 25% change or more significant change in activity following knockout of the perturbed node, and sensitivity is the number of nodes that will affect the target by a 25% change or greater when knocked out.

The CM proliferation signaling network was exported from Netflux into Cytoscape (Shannon et al., 2003) for topological analysis. The Network Analyzer (Assenov et al., 2008) plug-in was used to calculate the network’s topological properties. The correlation coefficient for matching topological to functional metrics was computed using the fitlim function in MATLAB. The functional metrics as defined by the sensitivity analysis were: (1) influence, the number of nodes with an activity change more significant than 25% with the knockdown of node n; (2) sensitivity, the number of nodes that change the activity of node n by more than 25% when knocked down.

### Robustness Analysis

Network robustness to variation in model parameters was tested using a validation threshold of 5% absolute change (Tan, 2017). For each parameter (Ymax, w, n, and EC50), new values were generated by sampling from a uniform random distribution with indicated half-width about the original parameter value. 100 new parameter sets were created for each distribution range for each parameter, and simulations were run to compare model predictions with literature observations. No changes in validation accuracy resulted from varying tau or Yinit.

### Materials

Williams E Medium (A1217601), cocktail B supplement (CM4000), Penicillin-Streptomycin(P/S), Alexa Fluor 568 antibody, AlexaFluor 488, AlexaFluor 680, and DAPI were purchased from Life Technologies. α-actinin antibody (A7811) was purchased from Sigma. Ki67 Monoclonal Antibody was purchased from Invitrogen. Pi3K inhibitor (Ly294002-S1105) was purchased from Selleck Chem. cMyc inhibitor (10058-F4(ab145065)) was purchased from Abcam. FoxM1 inhibitors, RCM1(6845), and FDI6(SML1392) were purchased from Tocris and Millipore Sigma. TT10 drug (2230640-94-3) was purchased from Aobius.

### Cell Culture

Cardiac myocytes were isolated from 1-2-day-old Sprague-Dawley rats using a Neomyt isolation kit (Cellutron, USA). The cells were cultured in plating media (low-glucose Dulbecco’s modified eagle media (DMEM), 17% M199, 10% horse serum, 5% fetal bovine serum (FBS), 1% L-Glutamine, 10 U/mL penicillin, and 50mg/mL streptomycin) at a density of 30,000 cells per well of a 96-well Corning CellBIND plate. After 48 hours of incubation with daily media change, the medium was replaced with WE+B medium and serum-starved for 4+hours before treatment.

### Proliferation Assay

Cells were treated with respective drugs in serum-free media for two days. DMSO (0.2%) and Nrg (50 ng/mL) were used as the negative and positive controls.

### Immunofluorescence and Imaging

After two days of treatment, the cells were fixed in 4% paraformaldehyde (PFA) and fluorescently labeled with DAPI, monoclonal anti -α-actinin or cardiac troponin T (cTnT), anti Ki67, and phospho-Histone 3 (pHH3). Cells were then imaged with the Operetta CLS high-content imaging system (Perkin-Elmer) using a 10x 0.3 NA objective.

### Image Processing

Image analysis scripts were developed in MATLAB to segment and classify the cells automatically. Nuclear segmentation methods adapted from Woo et al. and Bass et al. apply to images of nuclei stained with DAPI (Bass et al., 2012; Woo et al., 2019). Briefly, a median filter with a window size of three pixels was applied to smooth the images, and the blurred nuclei were segmented using an Otsu threshold. Next, we identified clumps of nuclei by measuring the circularity of the segment’s nuclei (<0.75). Objects with a low circularity factor were further separated by applying an erosion operation and then a watershed transform. Finally, objects outside the range of cardiomyocyte nuclei and those touching the image borders were removed. A mask of these objects was used to measure the integrated intensities of DAPI, α-actinin, EdU, Ki67, and pHH3. Thresholds for determining whether a nucleus was positive for a label were calculated as a fraction of the standard deviation from the model of the integrated intensities.

### Statistics

Primary rat cardiomyocytes were isolated from two different isolations. For each isolation, the treatment groups had at least 4 replicate wells for a total of 8 replicates per experimental group. We performed a two-way ANOVA on all experimental groups with a post-hoc Dunnett’s test comparing the two different isolations. There were 200-800 nuclei countered per well in the experiments. Error bars represent the standard error of the mean. Statistical significance was set at p < 0.05 or p < 0.01 as shown.

### mRNA sequencing analysis

RNA-seq data was retrieved from the GEO repository database (GSE123457) consisting of RNA extracted from Bead-bound PCM1+ nuclei from two control samples and two samples of YAP5SA overexpressing CMs (Monroe et al., 2019). Relative mRNA abundance values for Pi3Kca, Myc, and FoxM1 were averaged for the two control samples and the two YAP5SA overexpressing samples. Significance was estimated by performing a student’s t-test.

### Assay for Transpose Accessible Chromatin Sequencing Analysis

ATAC-seq data was retrieved from the GEO repository database (GSE123457) consisting of two control samples of cardiomyocyte (CM) nuclei extracted from a Myh6-MCM mouse strain and two YAP5SA overexpressing samples of CM nuclei extracted from a YAP5SA gain-of-function transgenic mouse strain (Monroe et al., 2019). Approximately 50,000 CM nuclei were extracted for each sample as input for ATAC-seq. Paired end 2×75 bp sequencing was performed using the Illumina Nextseq 500 instrument and reads were mapped to the mouse genome (mm10) using Bowtie2. We downloaded the processed BigWig files to use for peak-annotation at the Pik3ca, Myc, and Foxm1 loci. BigWig files were converted to bedgraph files using the UCSC-tools/3.7.4 (Kent et al., 2010) conversion package bigWigToBedGraph. Bedgraph files for each experimental condition were then merged using BEDTools unionbedg(Quinlan & Hall, 2010). Finally, we called the accessible chromatin region peaks for each of the two merged files using MACS2 bdgpeakcall (Zhang et al., 2008) with the default p-value cutoff threshold of 1e-5. Bedgraph and narrowPeak files were visualized using the Integrative Genomics Viewer (Robinson et al., 2011).

## Results

### A predictive computational model of the cardiomyocyte proliferation regulatory network

Although there has been substantial progress in discovering key regulators of cardiomyocyte proliferation, their interconnections have yet to be defined at a network level. For this reason, we manually constructed a regulatory network model using ∼23 literature articles that described general protein interactions that regulate proliferation. We also utilized resources such as Signor, which represents the cell cycle G1/S and G2/M phase transition pathways in humans (Licata et al., 2020); KEGG, which describes the mitotic cell cycle progression (*KEGG PATHWAY: Cell Cycle - Homo Sapiens (Human)*, n.d.); and Cell Signaling, which also describes cell cycle regulation pathways (*Cell Cycle Regulation Pathways*, n.d.). Literature descriptions of theseanal interactions and mechanisms were categorized based on whether they described a direct interaction (e.g. RB1 inhibits Cyclin D) or an indirect relationship (e.g. CDC25A overexpression enhances DNA replication). The former type was used for the model development category, while the latter was used later to validate the model. Literature articles used for model development came from multiple cell types due to limited CM data.

The CM proliferation signaling network integrates six modules that describe the interactions between signaling pathways and cell cycle regulators (**Figure 1A**). The network links regulatory modules of the cell cycle: core cyclin-dependent kinases (DNA replication module) regulating the G1/S checkpoint; mitosis and DNA damage responses regulating the G2/M checkpoint (mitosis module); as well as well-studied signaling pathways in the heart: growth factor and Hippo modules (Xin et al., 2011, 2013; Zheng et al., 2020). Each module contains proteins, transcription factors, or genes that were key to the overall process. For instance, the DNA replication module incorporates major cyclin and cyclin-CDK complexes and cyclin inhibitors that play a role in the cell cycle progression through the G1/S checkpoint (**Figure 1B**). The mitosis module incorporates DNA damage cues (ATM and ATR) that activate parallel pathways that inhibit the primary regulator of mitosis, cyclin B/CDK (**Figure 1C**). The main effectors of cytokinesis in the heart have not been well studied. Our cytokinesis module relies on nodes and interactions that have been primarily studied in other cell types. This included those examined in CM regulation, such as anillin, shown by Engel et al. as a critical process for cytokinesis in cardiomyocytes, and Ect2, shown to contribute to cytokinesis and increased binucleated cells (**Figure 1D**) (Bergmann, 2021; Engel et al., 2006; Jiang et al., 2019; Liu et al., 2019). Extracellular factors have been identified to activate intracellular receptors involved in cardiomyocyte proliferation: FGF1, Nrg1, and IGF1 (Rebouças et al., 2016). The growth factor module shows how these factors interlink with cell cycle progression regulators (**Figure 1E**). The Hippo signaling pathway is among the best-established modules for regulating CM proliferation. The Hippo module incorporates regulators such as YAP and TEAD that substantially affect cardiomyocyte proliferation (Xin et al., 2013) (**Figure 1F**). The output module interlinks the phenotypic outputs of the modules, usually used to signify proliferation in *in vivo* and *in vitro* experiments: DNA replication, often measured by EdU or Ki67; mitosis, phospho Histone 3 (pHH3); cytokinesis, Aurora B; binucleated and polyploid cells. For example, based on current knowledge of the organization of the cell cycle, in our model, activation of DNA replication and Cyclin B/CDK2 is required for mitosis. DNA replication can lead to polyploidization, but the activation of Cyclin B/CDK inhibits polyploidization (**Figure 1G**). Nodes or regulators with dashed borders that overlap between modules were used to connect the modules to create the CM proliferation network model. For example, cMyc is found in the Hippo, growth factor, and DNA replication modules, so those reactions were combined (**Figure S1**). **Supplementary Figure 1** shows the entire network connected with overlapping nodes, with the network activity simulated in the basal state.

**Figure 1:**
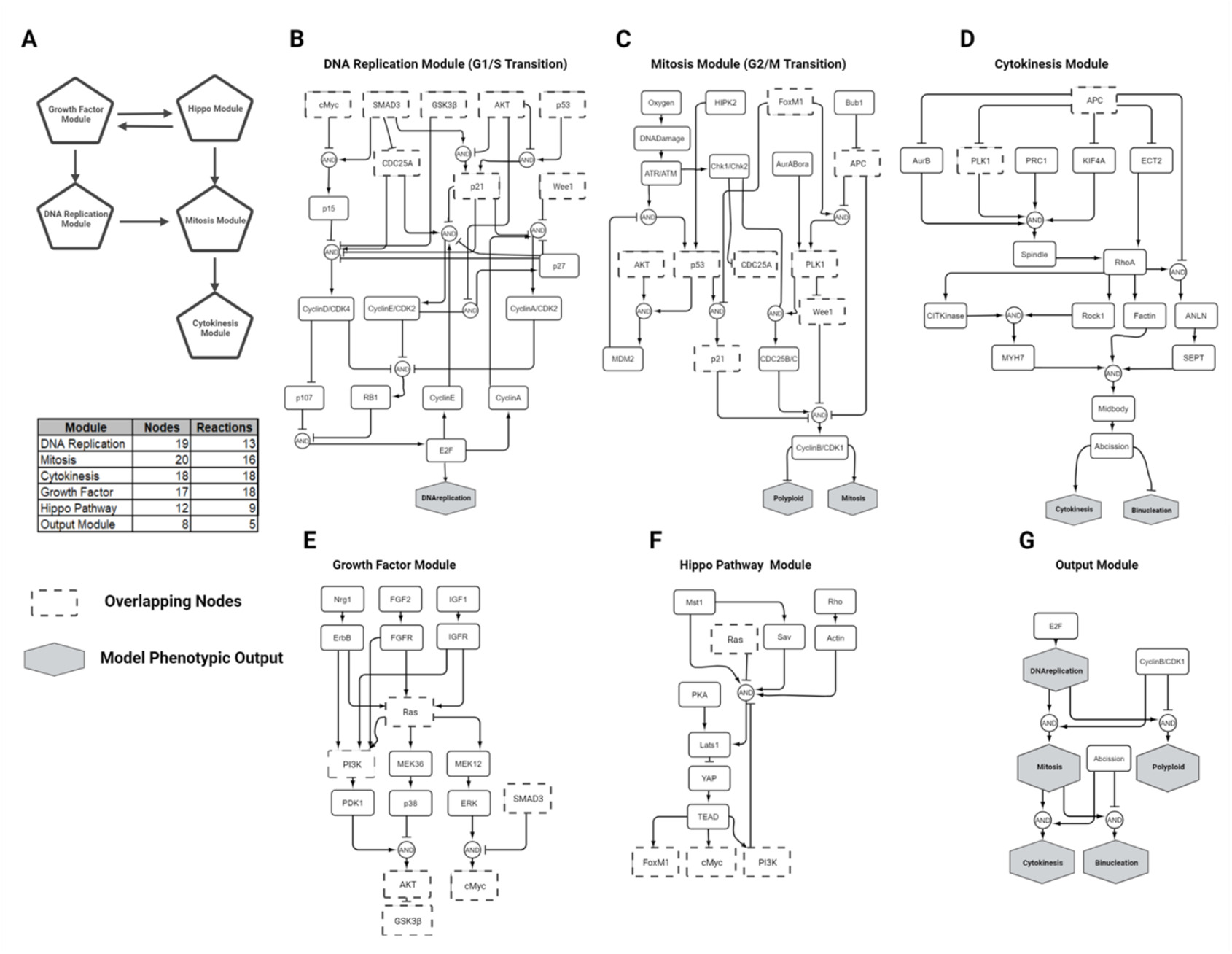
A computational model of the cardiomyocyte proliferation regulatory network. (A) Schematic that maps the model’s five regulatory modules and their interaction. The table describes the number of nodes and reactions within each module. (B-G) Activity flow diagram for each module, with dashed boxes representing those nodes that appear in multiple modules. Phenotypic outputs are represented as grey hexagons.

To convert the network into a predictive computational tool, network reactions were modeled using logic-based differential equations (LDEs) (Ryall et al., 2012; Tan, 2017; Zeigler et al., 2016). In this approach, the normalized activation of each node is represented by ordinary differential equations with saturating Hill functions and continuous logical AND or OR logic gates to characterize pathway crosstalk. OR gating is used when each input to a node is sufficient but not necessary for activation, whereas AND gating is used when each input is required. As in previous models (Tan, 2017; Zeigler et al., 2016), default values were used for network parameters. Preservation of network predictions based on these constraints has been previously demonstrated, although individual parameters can be tuned when necessary by fitting experimental measurements (Tan, 2017). Logic-based differential equations were generated in Netflux and implemented in MATLAB to predict model dynamics and simulate perturbations to the model parameters (**Supplementary Video**). Baseline conditions of model inputs were adjusted based on literature information about their presence in cardiomyocyte phenotypes (neonatal, adult, and embryonic validation). For instance, MST1, a major YAP/TAZ/Tead signaling inhibitor, is thought to have a high activity within the adult phenotype, so the node input was set between 0.8 and 1.

To examine the predictive accuracy of our model, we simulated activity changes in response to reported network perturbations. Experimental data were compared to model predictions using a 1% threshold for defining activity change. For example, the model predicted that adding growth factors Nrg and FGF with p38 inhibition would increase the percentage of cells going through DNA replication (**Figure 2A**). These predictions are consistent with the published data in cardiomyocytes from Engel et al., which found that p38 activity augments growth factor-mediated DNA synthesis in neonatal cardiomyocytes (Engel et al., 2005).

**Figure 2:**
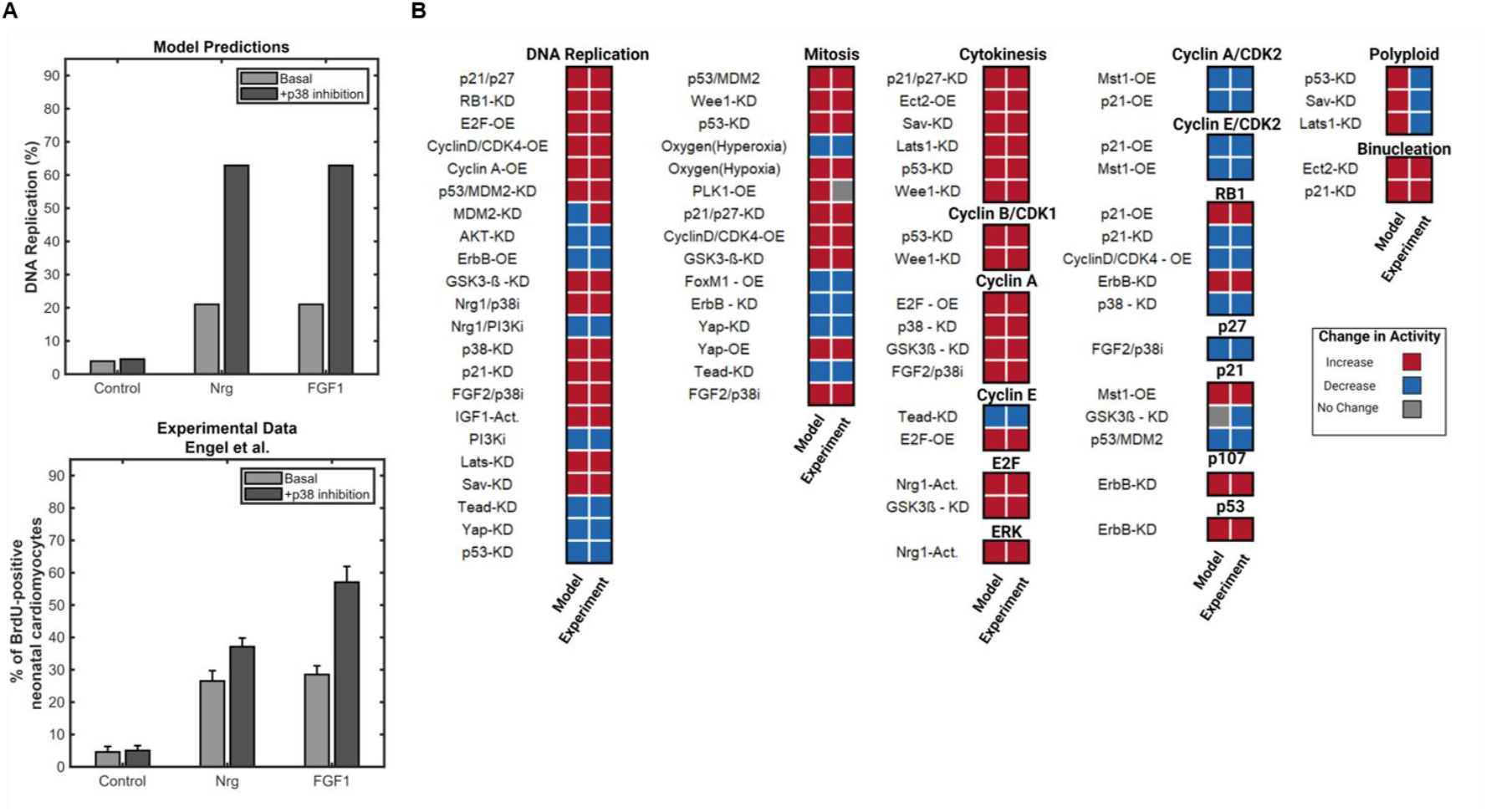
Validation of model predictions against experimental data from the literature. (A) Model prediction of DNA synthesis in response to p38 inhibition and a combination of p38 inhibition with growth factors, compared to experimental data from Engel et al. 2005 (Engel et al., 2005) (B) Qualitative comparison of model predictions in response to the noted stimuli, compared with published experimental observations of *in vitro* and *in vivo* cardiomyocyte proliferation. The model validates 72 of 76 (94.6%) experimental data not used to construct the model.

We compared these predictions with 76 published experimental observations of *in vivo* or *in vitro* cardiomyocytes. These observations included only those experiments performed using cardiomyocytes and were gathered from ∼33 literature articles not used for model construction. Based on statistics from the original studies, observations from the literature were encoded as increase, decrease, or no change. Overall, the model accurately predicts 72 of the 76 (94.7%) of the observations. **Figure 2B** summarizes the validation of three different validation relationship types: input-output, inhibition, and overexpression. We first focused on validating the phenotypic outputs of the model by searching for descriptions of experiments that measured: EdU incorporation or Ki67 expression (indication of DNA replication), pHH3 (indicating mitosis), and Aurora B at the midbody (cytokinesis). We included validations of individual nodes by looking for gene expression measurements in literature to expand our validation. We validated important cyclins and their CDK complexes (cyclin A, B, and E) as well as well-characterized cell cycle inhibitors (RB1, p27, p107, p21, and p53) that affect proliferation (**Figure 2B**). The model predicts 94.7% of observations found in the literature, but there are some exceptions. For example, the three model predictions and experimental observations for polyploidization did not match, indicating the complexity surrounding the possibility of cell phenotypes in cardiomyocyte populations.

As the true values of parameters affecting network node and weights are uncertain, we sampled parameters from uniform random distributions to evaluate the robustness of the model validation to the parameter values. Consistent with previous studies of other networks (Kraeutler et al., 2010; Tan, 2017; Zeigler et al., 2016), the validation accuracy is robust (>80%) to variation in model parameters over a uniform random distribution of up to 30% for ymax and EC50 and up to 30% or more for w (**Figure S4**). This shows that model predictions are robust despite parameter uncertainty.

### Influential drivers of CM proliferation with Nrg1 stimulation

After validating the model’s predictive capacity, we performed a network-wide sensitivity analysis to characterize the functional roles of the proteins and genes in the cardiomyocyte proliferation signaling network. As Nrg1 has previously been found to increase the proliferation of cardiomyocytes (Bersell et al., 2009; D’Uva et al., 2015; D’Uva & Tzahor, 2015; Rupert & Coulombe, 2015), we asked what nodes are most influential in driving Nrg1-dependent proliferation. We identified the most influential nodes as those whose knockdown produces the highest summed change in the phenotypic outputs of the network, along with the top measured nodes (**Figure 3A**). This analysis allows us to predict the network inhibition response of specific proteins, receptors, or genes. The top-most changed, or sensitive, nodes included key cell cycle regulatory proteins such as cyclins, cyclin-CDK complexes, and cell cycle inhibitory proteins. Additionally, we can see that the phenotypic outputs are highly influenced by the knockdown of these most influential nodes in the activated Nrg1 context.

**Figure 3:**
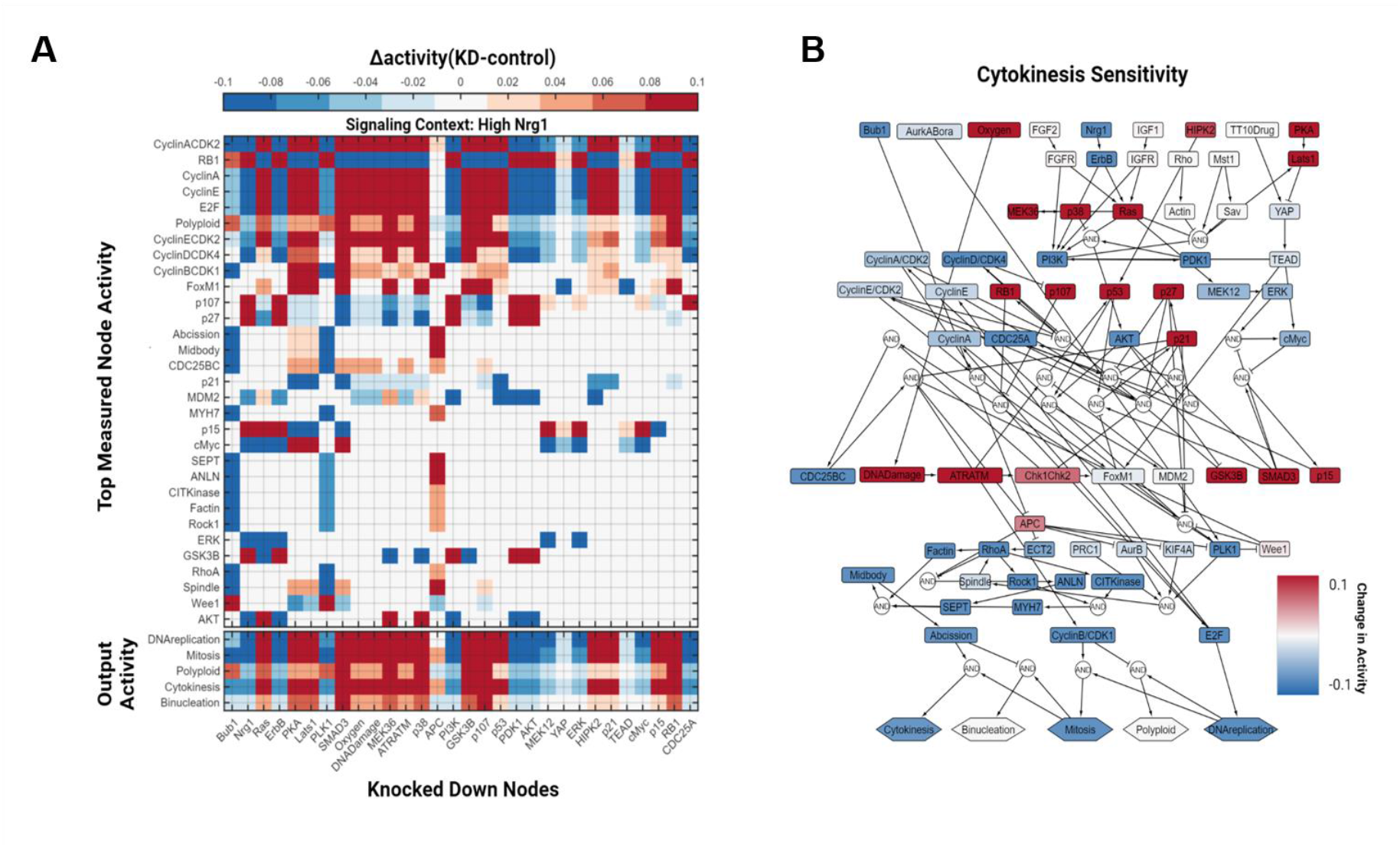
Virtual knockdown screen to predict regulators of CM proliferation with Nrg1 stimulation. (A) Knockdown of individual nodes (columns) in the context of high Nrg1 (input = 0.8) and the effect on network node activities (rows). (B) Network visualization of the effect of node knockdowns on predicted cytokinesis.

To further explore how the knockdown of the model nodes affects phenotypic outputs, we examined the network with a comprehensive virtual knockdown screen for regulators of cytokinesis. Cytokinesis requires the successful progression through the G1/S phase (DNA replication module) and the G2/M phase (mitosis module) (Auchampach et al., 2022; Milliron et al., 2019; Mohamed et al., 2018). In the model’s baseline state, only a few nodes (PKA, Lats1, RB1, and SMAD3) strongly influence cytokinesis activity, as cytokinesis is very low (**Figure S6A**). In contrast, the knockdown screen in the context of high Nrg1 (**Figure 3B**) shows nodes that are not only negative regulators of cytokinesis but important for DNA replication (e.g., RB1, p107, GSK3β, and other CKIs) and mitosis (e.g., ATR/ATM, APC, p53 and p21) regulation (see outputs in **Figure 3A**). In addition to negative regulators, there are positive regulators that strongly influence cytokinesis. These include nodes such as CyclinD/CDK4, a direct regulator of DNA replication, PI3K and AKT, which are important for both, and CDC25BC, an upstream regulator of mitosis. In the increased Nrg1 state, we see nodes that become important for cytokinesis from both DNA replication and mitosis modules.

### Context-dependent regulation of CM proliferation

Next, we asked whether the influence of nodes in the cardiomyocyte proliferation network differs depending on the proliferative stimulus. We compared the influence of individual nodes on the DNA replication, mitosis, and cytokinesis modules in the context of increased Nrg1, YAP activity, or baseline conditions. We first performed a sensitivity analysis under baseline conditions (**Figure S2**). We identified the 25 nodes under basal conditions with the highest influence over network activity and the phenotypic outputs. The basal state’s most influential nodes included those mediating signals from the growth factor and Hippo signaling module to the rest of the network: PI3K/AKT, GSK3B, SMAD3, Nrg1/ErbB, and Lats1 (**Figure S3A**).

We next visualized the network under YAP stimulation (**Figure S3C**). With an increase in activated YAP, several nodes in the DNA replication module change in influence compared basal state or Nrg1 context. For instance, with increased YAP activity AKT, CDC25A, and key cyclins are the most influential nodes, while cell cycle inhibitors are lowered in their influence (**Figure 4A**). Nodes that were highly influential with high Nrg1 decreased in influence with high YAP, while nodes that weren’t considered as influential in the Nrg1 state became highly influential in the high YAP state, such as CyclinA/CDK2, CDC25BC, and FOXM1. Overall, the most influential nodes in the mitosis and the cytokinesis modules stay relatively the same within each signaling context (**Figure 4A**). In contrast, the regulation of the DNA replication module is highly context-dependent.

**Figure 4:**
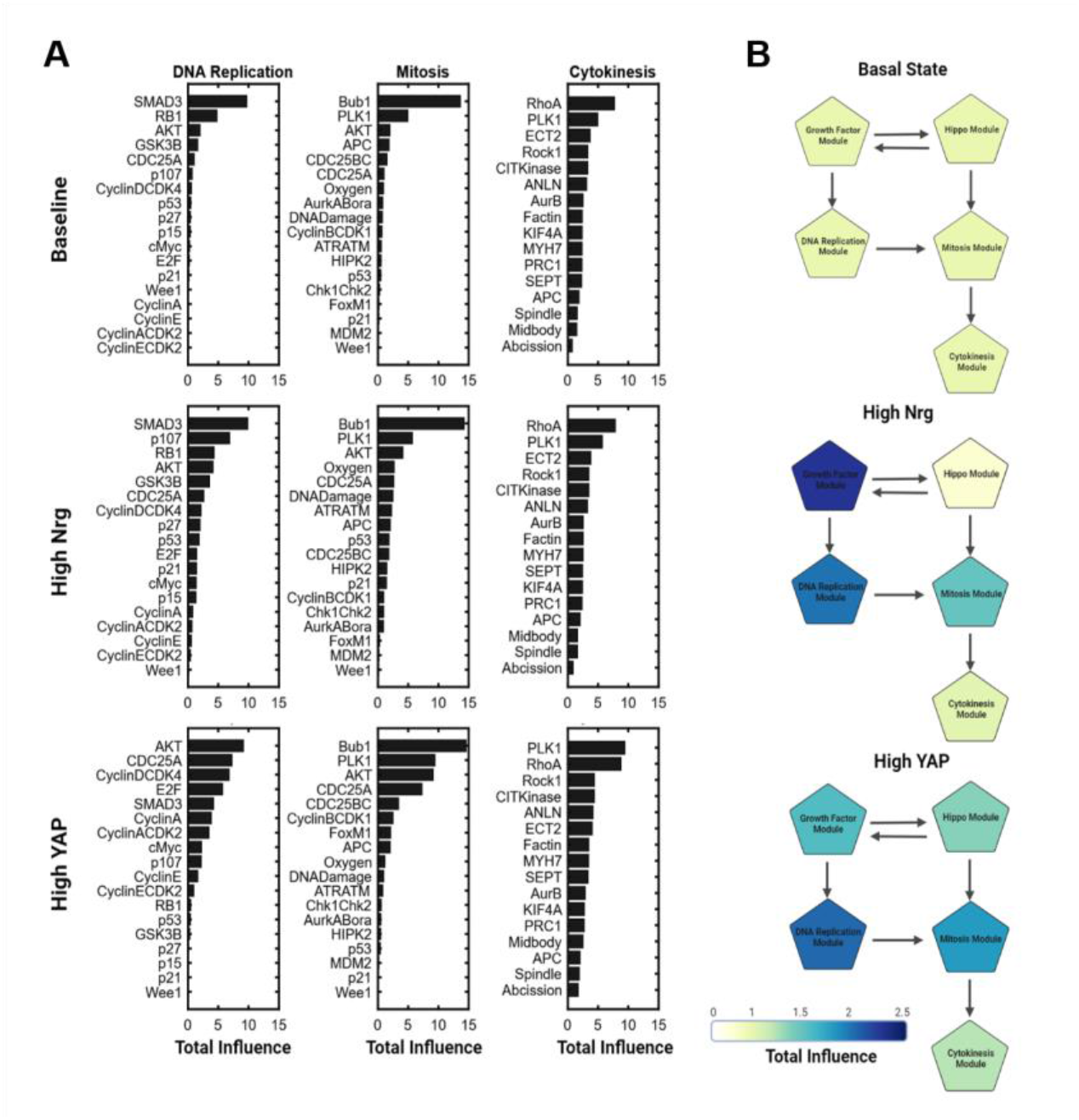
Influence of node knockdowns shifts with context, revealing crosstalk from Hippo to Growth Factor modules. (A) Total influence of node knockdowns on the DNA replication, mitosis, and cytokinesis modules, compared across multiple signaling contexts: baseline, high Nrg, and high YAP. Total influence sums the overall effect of a node knockdown on a network module. (B) The total influence of each network module varies depending on whether a basal state, high Nrg, or high YAP signaling context is applied.

To gain a more global view of the relationships between network modules, we computed the total influence of each module and quantified how the total influence of each module varies with the signaling context (**Figure 4B**). As expected, in the context of increased Nrg1, the growth factor module increases in influence, while the Hippo module slightly decreases compared to the basal state. This is followed by an increase in the influence of DNA replication, mitosis, and cytokinesis modules consistent with increased proliferation. In contrast, stimulation of YAP increases the influence of both the Hippo and growth factor modules, indicating underappreciated crosstalk between these modules. This leads to an even greater increase in the influence of DNA replication, mitosis, and cytokinesis module compared to the basal context or high Nrg1 context. Overall, these simulations identify crosstalk from YAP to the growth factor module that is predicted to be important for context-dependent CM proliferation.

### Crosstalk between modules of the CM regulatory network

The role of the Hippo pathway in cardiac regeneration and as a potential therapeutic target is well established (Wang et al., 2018; Zheng et al., 2020). To identify potential mediators of YAP signaling that crosstalk to the growth factor module, we simulated YAP stimulation and found nodes with greater than 15% change in activity (**Figure 5A**). About 70% of the DNA replication module had a significant increase in response to an increase in YAP activity and network outputs. Interestingly, FoxM1, PI3K, and cMyc almost double their activation with an increase in YAP, and all three are directly regulated by YAP’s target transcription factor TEAD in the model (**Figure 5B**).

**Figure 5:**
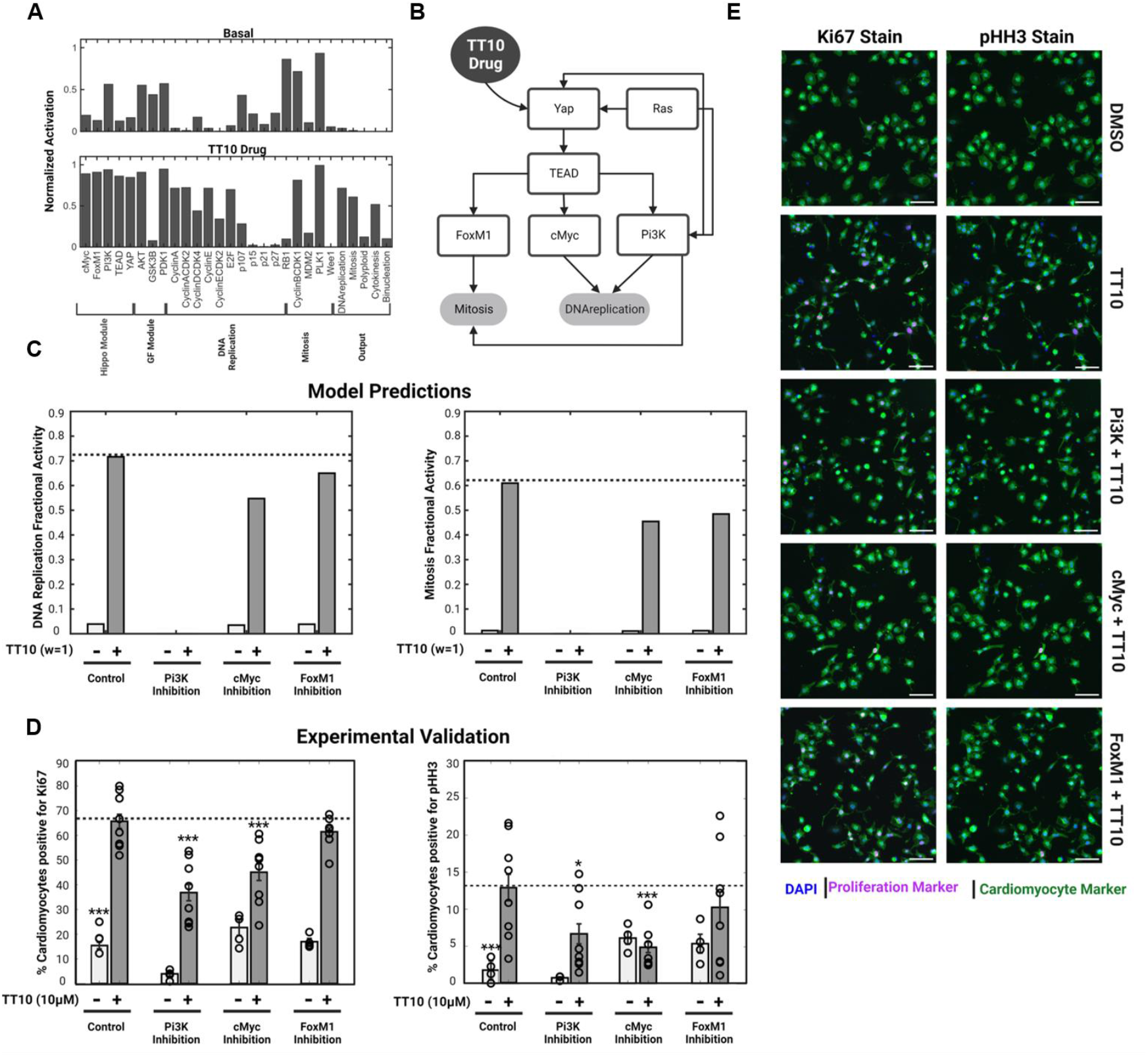
Prediction and experimental validation of cardiomyocyte proliferation mediated by the Hippo pathway via PI3K, cMyc, and FoxM1. (A) Network nodes affected by at least 15% in response to high YAP, grouped by module. (B) Network hypothesis of key hubs in the Hippo signaling pathway and how they affect the phenotypic outputs. (C) Quantification of model predictions of inhibitors for PI3K, cMyc, and FOXM1 alone and in the context of high YAP. (D) Quantification of Ki67 and pHH3 from experiments in rat neonatal CMs labeling Ki67 (cell cycle activity) or pHH3 (mitosis) in response to 10 μM TT-10, PI3K inhibitor (1 μM Ly294002), cMyc inhibitor (20 μM 10058-F4), or FOXM1 inhibitor (1 μM RCM1). (E) Representative images from the experiment are described in (D).

We further explored whether PI3K, FoxM1, and cMyc mediate the proliferative effects of YAP. Previously, Lin et al. found in neonatal rat ventricular cardiomyocytes that YAP functions directly upstream of PI3K to enhance its expression (Lin et al., 2015). In addition, they found that Pik3cb is a crucial direct target for YAP that links the Hippo-YAP pathway to the PI3K-AKT signaling pathway and promotes CM proliferation and survival (Lin et al., 2015)]. Other systems have shown the connection between YAP and FoxM1 in malignant mesothelioma cells, showing that YAP regulates FoxM1 transcription directly through TEAD (Mizuno et al., 2012).

We simulated the addition of TT10, a YAP-activating drug (Hara et al., 2018), coupled with inhibition of PI3K, FoxM1, or cMyc, on CM proliferation. As in past experiments, the model predicted that activating YAP with TT10 greatly increases the percentage of cells in the DNA replication and mitosis phases (Hara et al., 2018). Further, the model predicted that inhibition of PI3K, FoxM1, or cMyc would reduce the proliferative effect of TT10. To test these predictions experimentally, we cultured neonatal rat cardiomyocytes with combined activation of YAP (10 μM TT-10) and inhibition of PI3K (1 μM Ly294002), cMyc (100 μM 10058-F4), or FoxM1 (1 μM RCM1). As shown in Figure 5D, the increased cell cycle activity (Ki67) and mitosis (pHH3) induced by YAP activator TT10 were attenuated by inhibition of cMyc or PI3K but not FoxM1. These neonatal *in vitro* results validate the model prediction that YAP-induced proliferation is mediated by module crosstalk involving cMyc and PI3K.

We next asked if YAP may regulate the predicted targets of PI3K, cMyc, and FOXM1 in vivo. In Monroe et al. (Monroe et al., 2019), an active version of YAP, termed YAP5SA, was overexpressed in cardiomyocytes in adult mice. To examine whether YAP induces the expression of PI3K, cMyc, and FOXM1 in vivo, we analyzed RNA-seq data from Monroe et al. (Monroe et al., 2019). Indeed, cardiac overexpression of YAP5SA increased mRNA for Myc and Foxm1 (**Figure 6B**). To examine whether there is coincident remodeling of the chromatin within the promoter regions of these genes, we then analyzed ATAC-seq data (Monroe et al., 2019) from the same mouse model of YAPS5A (**Figure 6C-6E**). Promoters of both Myc and PI3Kca exhibited an altered chromatin open state. Only cMyc demonstrated fully consistent effects on YAP-dependent CM proliferation, YAP-dependent mRNA, and YAP-dependent chromatin opening, as predicted by the model.

**Figure 6:**
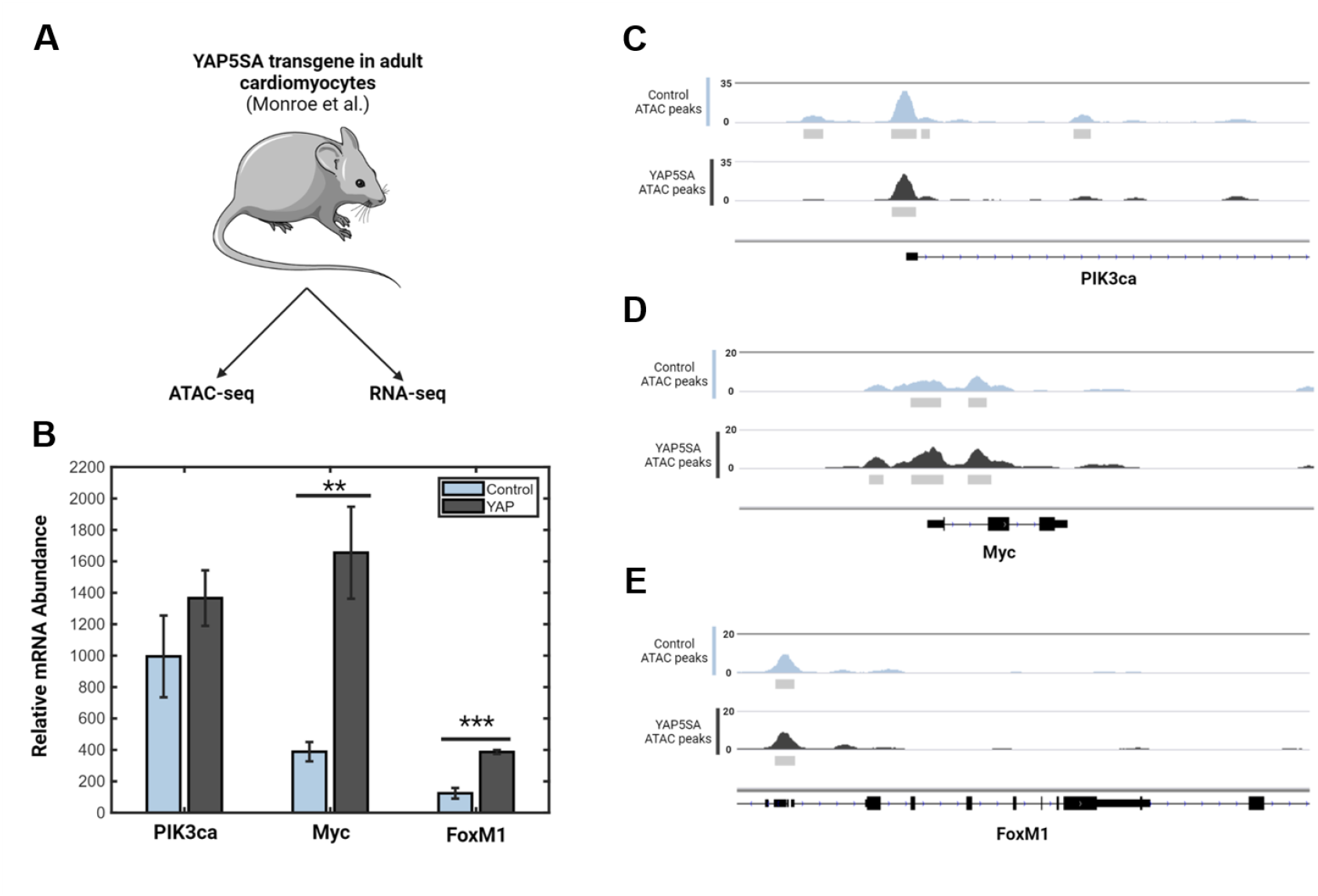
In vivo RNA-seq and ATAC-seq further validate the model-predicted role of PI3K, cMyc, and FoxM1 in YAP regulation of CM proliferation. (A) Schematic of experimental design of cardiac constitutively active YAP5SA transgenic mice (Monroe et al., 2019). (B) RNA-seq data showing expression of genes with and without the YAP5SA transgene. (C-E) ATAC-seq data describing open chromatin remodeling between genes with and without the YAP5SA transgene. Grey bars denote statistically significant regions of accessible chromatin (p<0.00001). Scale bars represent the genome browser track height and are proportional to the number of reads mapped to each genomic position.

## Discussion

In this study, we developed a dynamic map of the cardiomyocyte proliferation network to identify molecular drivers and network principles of CM proliferation. By creating and validating a predictive model of the cardiomyocyte proliferation network, we showed how the crosstalk between signaling pathways and core cell cycle regulators allows cardiomyocytes to progress through key cell cycle checkpoints (DNA replication, mitosis, and cytokinesis). Our model integrates five key modules that define cardiomyocyte proliferation and 72 CM proliferation nodes connected by 86 reactions. The model was validated at 94% compared to independent, published experimental studies in cardiomyocytes. A sensitivity analysis of the model revealed the context dependence of the cell cycle modules. Network analysis identified key hubs of the Hippo signaling pathway that crosstalk with the growth factor module (cMyc, PI3K, and FoxM1), and these hubs were experimentally validated in neonatal rat cardiomyocytes. We further validated that cardiac-specific expression of active YAP regulates mRNA and chromatin opening of Myc in vivo.

### Identification of knowledge gaps in cardiomyocyte proliferation

While the model validated 94% of input-output relationships from observations not used in model creation, five input-output relationships were incorrectly predicted. For instance, Mdm2 (murine double minute 2), which targets p53 (Figure 1C), was shown by Stanley-Hasnain et al. to increase DNA synthesis in cardiomyocytes when knocked down (Stanley-Hasnain et al., 2017). However, our model predicted that it decreased DNA synthesis activity. MDM2 plays a diverse role in the cardiovascular systems, and studies to elucidate its interaction with upstream and downstream regulators are needed (Hauck et al., 2017; Lam & Roudier, 2019; Stanley-Hasnain et al., 2017). The majority of the incorrect predictions involved output responses for the polyploid phenotype. For example, the model predicted a decrease in polyploid cells in response to Lats1 knockdown when experimental data from literature showed an increase (Heallen et al., 2013). In a review by Derks and Bergmann, they find that despite numerous explanations for cardiomyocyte polyploidization that have been proposed, more work needs to be done to identify distinct mechanisms regulating cardiomyocyte ploidy (Derks & Bergmann, 2020). Similar findings were reported in Kirillova et al. (Kirillova et al., 2021). These incorrect predictions highlight areas for which future model revision and experiments would be necessary.

### Network crosstalk and its control of cardiomyocyte proliferation

This predictive model provides a framework to identify the most effective way to induce CM proliferation in response to specific stimuli. Neuregulin-1 (Nrg1) is well described to promote cardiomyocyte proliferation, and is a prime molecular target to promote cardiac regeneration. In addition, YAP, a major effector of Hippo, is essential in regulating cardiomyocyte proliferation and survival (Del Re et al., 2013; Heallen et al., 2011, 2013; Lin et al., 2015; von Gise et al., 2012; Xin et al., 2013). Using the influence metric from our sensitivity analysis, we compared total module influence in the basal, high Nrg1, and high YAP contexts. Analysis of our model identified crosstalk between the growth factors and Hippo signaling modules. When YAP is increased, not only does the Hippo module itself increase, but the influence of the growth factor module also increases. For this reason, we investigated the role of YAP in the downstream regulation of cardiomyocyte proliferation.

Previous studies of YAP signaling focused on derepressing upstream inhibitors of YAP to increase proliferation (Flinn et al., 2020). More specifically, targeting of MST1/2, SAV, and Lats 1 to activate YAP (Heallen et al., 2011, 2013; Monroe et al., 2019). However, the direct targets of activated YAP that control downstream regulation are unclear. Our model predicted three main potential hubs (PI3K, cMyc, and Foxm1) connecting YAP to the growth factor and mitosis modules. We tested whether the knockdown of these nodes would attenuate the proliferative effects of increased YAP. Analyzing both model predictions and experimental validations, we found that the inhibition of PI3K and cMyc abrogates the proliferative effect of increasing YAP activity. A study by Mizuno et al. showed that YAP induced proliferation by upregulation of FoxM1 and other cell cycle genes in malignant mesothelioma (Mizuno et al., 2012). In addition, Bolte et al. implicated FoxM1 as necessary for myocardial development (Bolte et al., 2011). Others have connected YAP to an increase in the expression of PI3K (Lin et al., 2015). To our knowledge, the relationship between YAP, these three regulators, and their potential crosstalk to regulate cardiomyocyte proliferation has not been previously shown in perturbation experiments. Therefore, we tested whether the inhibition of these nodes would attenuate the effect of increasing YAP activity. Experimentally, we found that the inhibition of PI3K or cMyc abrogates the effect of increasing YAP, validating the model’s prediction. Analysis of RNA-seq data from YAP5SA expressing CMs from Monroe et al. (Monroe et al., 2019) showed a significant increase in Myc and FoxM1 expression but not PIK3ca. In addition, ATAC-seq showed changes in chromatin accessibility between control and YAP5SA overexpression in PIK3ca and Myc but not in FoxM1. This validates YAP to cMyc crosstalk as important for CM proliferation.

### Limitations and future directions

The construction of this network used logic-based differential equations and utilized default model parameters (but see **Figure S4**), which we have previously shown to exhibit a strong predictive accuracy for other networks (Kraeutler et al., 2010; Tan, 2017; Zeigler et al., 2016). As more quantitative data becomes available, we can adjust parameters to replicate different phenotypes or developmental stages, including the adult CM. For instance, in this study, 80% of the validations were from neonatal CMs, with less than 10% from adult CMs. The network model identifies gaps in the current understanding of cardiomyocyte proliferation signaling. Even though the five phenotypic outputs of the model have been extensively studied and verified by experimental data, the potential regulators that lead to binucleated or polyploid cells are still unclear. As are direct regulators of cytokinesis. Our model structure focuses on the current information on cell cycle signaling networks that meets specified criteria for inclusion in a cardiomyocyte-specific model. This model provides an initial network framework for integrating future discoveries in cardiomyocyte proliferation.

## Conclusion

We developed a predictive model of the cardiomyocyte proliferation signaling network that identifies the nodes and network structures that regulate the proliferation of cardiomyocytes. A sensitivity analysis of our model identified drivers of cardiomyocyte proliferation and showed that the drivers for DNA replication vary in influence with changes in signaling contexts. In contrast, node influence within mitosis and cytokinesis modules was more robust to context. Based on total module influence in different signaling contexts, the model predicted crosstalk from the Hippo module to the growth factor module involving cMyc, PI3K and FoxM1, with YAP to cMyc crosstalk validated experimentally and *in vivo*.

## Acknowledgements

We thank Dr. Mohammad Fallahi-Sichani for use of his Operetta CLS high-content imaging system.

## References

Assenov, Y., Ramírez, F., Schelhorn, S.-E., Lengauer, T., & Albrecht, M. (2008). Computing topological parameters of biological networks. Bioinformatics (Oxford, England), 24(2), 282–284. https://doi.org/10.1093/bioinformatics/btm554

Auchampach, J., Han, L., Huang, G. N., Kühn, B., Lough, J. W., O’Meara, C. C., Payumo, A. Y., Rosenthal, N. A., Sucov, H. M., Yutzey, K. E., & Patterson, M. (2022). Measuring cardiomyocyte cell-cycle activity and proliferation in the age of heart regeneration. American Journal of Physiology-Heart and Circulatory Physiology, 322(4), H579–H596. https://doi.org/10.1152/ajpheart.00666.2021

Bass, G. T., Ryall, K. A., Katikapalli, A., Taylor, B. E., Dang, S. T., Acton, S. T., & Saucerman, J. J. (2012). Automated image analysis identifies signaling pathways regulating distinct signatures of cardiac myocyte hypertrophy. Journal of Molecular and Cellular Cardiology, 52(5), 923–930. https://doi.org/10.1016/j.yjmcc.2011.11.009

Bergmann, O. (2021). Cardiomyocytes in congenital heart disease: Overcoming cytokinesis failure in tetralogy of Fallot. The Journal of Thoracic and Cardiovascular Surgery, 161(5), 1587–1590. https://doi.org/10.1016/j.jtcvs.2020.05.091

Bersell, K., Arab, S., Haring, B., & Kühn, B. (2009). Neuregulin1/ErbB4 signaling induces cardiomyocyte proliferation and repair of heart injury. Cell, 138(2), 257–270. https://doi.org/10.1016/j.cell.2009.04.060

Bertero, A., & Murry, C. E. (2018). Hallmarks of cardiac regeneration. Nature Reviews Cardiology, 15, 579–580. https://doi.org/10.1038/s41569-018-0079-8

Bolte, C., Zhang, Y., Wang, I.-C., Kalin, T. V., Molkentin, J. D., & Kalinichenko, V. V. (2011). Expression of Foxm1 Transcription Factor in Cardiomyocytes Is Required for Myocardial Development. PLoS ONE, 6(7), e22217. https://doi.org/10.1371/journal.pone.0022217

CDC. (2022, July 15). Heart Disease Facts | cdc.gov. Centers for Disease Control and Prevention. https://www.cdc.gov/heartdisease/facts.htm

Cell Cycle Regulation Pathways. (n.d.). Cell Signaling Technology. Retrieved June 15, 2022, from https://www.cellsignal.com/pathways/by-research/cell-cycle-regulation-pathways

Del Re, D. P., Yang, Y., Nakano, N., Cho, J., Zhai, P., Yamamoto, T., Zhang, N., Yabuta, N., Nojima, H., Pan, D., & Sadoshima, J. (2013). Yes-associated protein isoform 1 (Yap1) promotes cardiomyocyte survival and growth to protect against myocardial ischemic injury. The Journal of Biological Chemistry, 288(6), 3977–3988. https://doi.org/10.1074/jbc.M112.436311

Derks, W., & Bergmann, O. (2020). Polyploidy in Cardiomyocytes. Circulation Research, 126(4), 552–565. https://doi.org/10.1161/CIRCRESAHA.119.315408

D’Uva, G., Aharonov, A., Lauriola, M., Kain, D., Yahalom-Ronen, Y., Carvalho, S., Weisinger, K., Bassat, E., Rajchman, D., Yifa, O., Lysenko, M., Konfino, T., Hegesh, J., Brenner, O., Neeman, M., Yarden, Y., Leor, J., Sarig, R., Harvey, R. P., & Tzahor, E. (2015). ERBB2 triggers mammalian heart regeneration by promoting cardiomyocyte dedifferentiation and proliferation. Nature Cell Biology, 17(5), 627–638. https://doi.org/10.1038/ncb3149

D’Uva, G., & Tzahor, E. (2015). The key roles of ERBB2 in cardiac regeneration. Cell Cycle (Georgetown, Tex.), 14(15), 2383–2384. https://doi.org/10.1080/15384101.2015.1063292

Engel, F. B., Schebesta, M., Duong, M. T., Lu, G., Ren, S., Madwed, J. B., Jiang, H., Wang, Y., & Keating, M. T. (2005). P38 MAP kinase inhibition enables the proliferation of adult mammalian cardiomyocytes. Genes & Development, 19(10), 1175–1187. https://doi.org/10.1101/gad.1306705

Engel, F. B., Schebesta, M., & Keating, M. T. (2006). Anillin localization defect in cardiomyocyte binucleation. Journal of Molecular and Cellular Cardiology, 41(4), 601–612. https://doi.org/10.1016/j.yjmcc.2006.06.012

Eschenhagen, T., Bolli, R., & Braun, T. (2017). Cardiomyocyte Regeneration: A Consensus Statement. Circulation, 136(7), 680–686. https://doi.org/10.1161/CIRCULATIONAHA.117.029343

Eulalio, A., Mano, M., Ferro, M. D., Zentilin, L., Sinagra, G., Zacchigna, S., & Giacca, M. (2012). Functional screening identifies miRNAs inducing cardiac regeneration. Nature, 492(7429), 376–381. https://doi.org/10.1038/nature11739

Flinn, M. A., Link, B. A., & O’Meara, C. C. (2020). Upstream regulation of the Hippo-Yap pathway in cardiomyocyte regeneration. Seminars in Cell & Developmental Biology, 100, 11–19. https://doi.org/10.1016/j.semcdb.2019.09.004

Hara, H., Takeda, N., Kondo, M., Kubota, M., Saito, T., Maruyama, J., Fujiwara, T., Maemura, S., Ito, M., Naito, A. T., Harada, M., Toko, H., Nomura, S., Kumagai, H., Ikeda, Y., Ueno, H., Takimoto, E., Akazawa, H., Morita, H., … Komuro, I. (2018). Discovery of a Small Molecule to Increase Cardiomyocytes and Protect the Heart After Ischemic Injury. JACC: Basic to Translational Science, 3(5), 639–653. https://doi.org/10.1016/j.jacbts.2018.07.005

Hashimoto, H., Olson, E. N., & Bassel-Duby, R. (2018). Therapeutic Approaches for Cardiac Regeneration and Repair. Nature Reviews Cardiology, 15(10), 585–600. https://doi.org/10.1038/s41569-018-0036-6

Hashmi, S., & Ahmad, H. R. (2019). Molecular switch model for cardiomyocyte proliferation. Cell Regeneration, 8(1), 12–20. https://doi.org/10.1016/j.cr.2018.11.002

Hauck, L., Stanley-Hasnain, S., Fung, A., Grothe, D., Rao, V., Mak, T. W., & Billia, F. (2017). Cardiac-specific ablation of the E3 ubiquitin ligase Mdm2 leads to oxidative stress, broad mitochondrial deficiency and early death. PLOS ONE, 12(12), e0189861. https://doi.org/10.1371/journal.pone.0189861

Heallen, T., Morikawa, Y., Leach, J., Tao, G., Willerson, J. T., Johnson, R. L., & Martin, J. F. (2013). Hippo signaling impedes adult heart regeneration. Development (Cambridge, England), 140(23), 4683–4690. https://doi.org/10.1242/dev.102798

Heallen, T., Zhang, M., Wang, J., Bonilla-Claudio, M., Klysik, E., Johnson, R. L., & Martin, J. F. (2011). Hippo Pathway Inhibits Wnt Signaling to Restrain Cardiomyocyte Proliferation and Heart Size. Science (New York, N.Y.), 332(6028), 458–461. https://doi.org/10.1126/science.1199010

Jiang, Y.-H., Zhu, Y., Chen, S., Wang, H.-L., Zhou, Y., Tang, F.-Q., Jian, Z., & Xiao, Y.-B. (2019). Re-enforcing hypoxia-induced polyploid cardiomyocytes enter cytokinesis through activation of β-catenin. Scientific Reports, 9, 17865. https://doi.org/10.1038/s41598-019-54334-4

KEGG PATHWAY: Cell cycle—Homo sapiens (human). (n.d.). Retrieved June 15, 2022, from https://www.genome.jp/pathway/hsa04110

Kent, W. J., Zweig, A. S., Barber, G., Hinrichs, A. S., & Karolchik, D. (2010). BigWig and BigBed: Enabling browsing of large distributed datasets. Bioinformatics (Oxford, England), 26(17), 2204–2207. https://doi.org/10.1093/bioinformatics/btq351

Kirillova, A., Han, L., Liu, H., & Kühn, B. (2021). Polyploid cardiomyocytes: Implications for heart regeneration. Development (Cambridge, England), 148(14), dev199401. https://doi.org/10.1242/dev.199401

Kraeutler, M. J., Soltis, A. R., & Saucerman, J. J. (2010). Modeling cardiac β-adrenergic signaling with normalized-Hill differential equations: Comparison with a biochemical model. BMC Systems Biology, 4, 157. https://doi.org/10.1186/1752-0509-4-157

Lam, B., & Roudier, E. (2019). Considering the Role of Murine Double Minute 2 in the Cardiovascular System? Frontiers in Cell and Developmental Biology, 7. https://www.frontiersin.org/articles/10.3389/fcell.2019.00320

Leach, J. P., & Martin, J. F. (2018). Cardiomyocyte Proliferation for Therapeutic Regeneration. Current Cardiology Reports, 20(63). https://doi.org/10.1007/s11886-018-1011-x

Licata, L., Lo Surdo, P., Iannuccelli, M., Palma, A., Micarelli, E., Perfetto, L., Peluso, D., Calderone, A., Castagnoli, L., & Cesareni, G. (2020). SIGNOR 2.0, the SIGnaling Network Open Resource 2.0: 2019 update. Nucleic Acids Research, 48(D1), D504–D510. https://doi.org/10.1093/nar/gkz949

Lin, Z., Zhou, P., von Gise, A., Gu, F., Ma, Q., Chen, J., Guo, H., van Gorp, P. R. R., Wang, D.-Z., & Pu, W. T. (2015). Pi3kcb Links Hippo-YAP and PI3K-AKT Signaling Pathways to Promote Cardiomyocyte Proliferation and Survival. Circulation Research, 116(1), 35–45. https://doi.org/10.1161/CIRCRESAHA.115.304457

Liu, H., Zhang, C.-H., Ammanamanchi, N., Suresh, S., Lewarchik, C., Rao, K., Uys, G. M., Han, L., Abrial, M., Yimlamai, D., Ganapathy, B., Guillermier, C., Chen, N., Khaladkar, M., Spaethling, J., Eberwine, J. H., Kim, J., Walsh, S., Choudhury, S., … Kühn, B. (2019). Control of cytokinesis by β-adrenergic receptors indicates an approach for regulating cardiomyocyte endowment. Science Translational Medicine, 11(513), eaaw6419. https://doi.org/10.1126/scitranslmed.aaw6419

Mann, D., & Bristow, M. (2005). Mechanisms and Models in Heart Failure. Circulation, 111(21), 2837–2849. https://doi.org/10.1161/circulationaha.104.500546

Milliron, H.-Y. Y., Weiland, M. J., Kort, E. J., & Jovinge, S. (2019). Isolation of Cardiomyocytes Undergoing Mitosis With Complete Cytokinesis. Circulation Research, 125(12), 1070–1086. https://doi.org/10.1161/CIRCRESAHA.119.314908

Mizuno, T., Murakami, H., Fujii, M., Ishiguro, F., Tanaka, I., Kondo, Y., Akatsuka, S., Toyokuni, S., Yokoi, K., Osada, H., & Sekido, Y. (2012). YAP induces malignant mesothelioma cell proliferation by upregulating transcription of cell cycle-promoting genes. Oncogene, 31(49), 5117–5122. https://doi.org/10.1038/onc.2012.5

Mohamed, T. M. A., Ang, Y.-S., Radzinsky, E., Zhou, P., Huang, Y., Elfenbein, A., Foley, A., Magnitsky, S., & Srivastava, D. (2018). Regulation of Cell Cycle to Stimulate Adult Cardiomyocyte Proliferation and Cardiac Regeneration. Cell, 173(1), 104-116.e12. https://doi.org/10.1016/j.cell.2018.02.014

Monroe, T. O., Hill, M. C., Morikawa, Y., Leach, J. P., Heallen, T., Cao, S., Krijger, P. H. L., de Laat, W., Wehrens, X. H. T., Rodney, G. G., & Martin, J. F. (2019). YAP Partially Reprograms Chromatin Accessibility to Directly Induce Adult Cardiogenesis In Vivo. Developmental Cell, 48(6), 765-779.e7. https://doi.org/10.1016/j.devcel.2019.01.017

Ouyang, Z., & Wei, K. (2021). MiRNA in cardiac development and regeneration. Cell Regeneration, 10, 14. https://doi.org/10.1186/s13619-021-00077-5

Oyama, K., El-Nachef, D., & MacLellan, W. (2013). Regeneration potential of adult cardiac myocytes. Cell Research, 23(8), 978–979. https://doi.org/10.1038/cr.2013.78

Payan, S. M., Hubert, F., & Rochais, F. (2020). Cardiomyocyte proliferation, a target for cardiac regeneration. Biochimica et Biophysica Acta (BBA) - Molecular Cell Research, 1867(3), 118461. https://doi.org/10.1016/j.bbamcr.2019.03.008

Quinlan, A. R., & Hall, I. M. (2010). BEDTools: A flexible suite of utilities for comparing genomic features. Bioinformatics (Oxford, England), 26(6), 841–842. https://doi.org/10.1093/bioinformatics/btq033

Rebouças, J. de S., Santos-Magalhães, N. S., & Formiga, F. R. (2016). Cardiac Regeneration using Growth Factors: Advances and Challenges. Arquivos Brasileiros de Cardiologia, 107(3), 271–275. https://doi.org/10.5935/abc.20160097

Robinson, J. T., Thorvaldsdóttir, H., Winckler, W., Guttman, M., Lander, E. S., Getz, G., & Mesirov, J. P. (2011). Integrative genomics viewer. Nature Biotechnology, 29(1), 24–26. https://doi.org/10.1038/nbt.1754

Rupert, C. E., & Coulombe, K. L. (2015). The Roles of Neuregulin-1 in Cardiac Development, Homeostasis, and Disease. Biomarker Insights, 10(Suppl 1), 1–9. https://doi.org/10.4137/BMI.S20061

Ryall, K. A., Holland, D. O., Delaney, K. A., Kraeutler, M. J., Parker, A. J., & Saucerman, J. J. (2012). Network reconstruction and systems analysis of cardiac myocyte hypertrophy signaling. The Journal of Biological Chemistry, 287(50), 42259–42268. https://doi.org/10.1074/jbc.M112.382937

Shannon, P., Markiel, A., Ozier, O., Baliga, N. S., Wang, J. T., Ramage, D., Amin, N., Schwikowski, B., & Ideker, T. (2003). Cytoscape: A software environment for integrated models of biomolecular interaction networks. Genome Research, 13(11), 2498–2504. https://doi.org/10.1101/gr.1239303

Stanley-Hasnain, S., Hauck, L., Grothe, D., Aschar-Sobbi, R., Beca, S., Butany, J., Backx, P. H., Mak, T. W., & Billia, F. (2017). P53 and Mdm2 act synergistically to maintain cardiac homeostasis and mediate cardiomyocyte cell cycle arrest through a network of microRNAs. Cell Cycle, 16(17), 1585–1600. https://doi.org/10.1080/15384101.2017.1346758

Tan, P. (2017). Predictive model identifies key network regulators of cardiomyocyte mechanics-signaling. PLOS Computational Biollogy, 13(11). https://doi.org/10.1371/journal.pcbi.1005854

Tian, Y., Liu, Y., Wang, T., Zhou, N., Kong, J., Chen, L., Snitow, M., Morley, M., Li, D., Petrenko, N., Zhou, S., Lu, M., Gao, E., Koch, W. J., Stewart, K. M., & Morrisey, E. E. (2015). A microRNA-Hippo pathway that promotes cardiomyocyte proliferation and cardiac regeneration in mice. Science Translational Medicine, 7(279), 279ra38. https://doi.org/10.1126/scitranslmed.3010841

Torrini, C., Cubero, R. J., Dirkx, E., Braga, L., Ali, H., Prosdocimo, G., Gutierrez, M. I., Collesi, C., Licastro, D., Zentilin, L., Mano, M., Zacchigna, S., Vendruscolo, M., Marsili, M., Samal, A., & Giacca, M. (2019). Common Regulatory Pathways Mediate Activity of MicroRNAs Inducing Cardiomyocyte Proliferation. Cell Reports, 27(9), 2759-2771.e5. https://doi.org/10.1016/j.celrep.2019.05.005

von Gise, A., Lin, Z., Schlegelmilch, K., Honor, L. B., Pan, G. M., Buck, J. N., Ma, Q., Ishiwata, T., Zhou, B., Camargo, F. D., & Pu, W. T. (2012). YAP1, the nuclear target of Hippo signaling, stimulates heart growth through cardiomyocyte proliferation but not hypertrophy. Proceedings of the National Academy of Sciences of the United States of America, 109(7), 2394–2399. https://doi.org/10.1073/pnas.1116136109

Wang, J., Liu, S., Heallen, T., & Martin, J. F. (2018). The Hippo pathway in the heart: Pivotal roles in development, disease, and regeneration. Nature Reviews Cardiology, 15(11), 672–684. https://doi.org/10.1038/s41569-018-0063-3

Woo, L. A., Tkachenko, S., Ding, M., Plowright, A. T., Engkvist, O., Andersson, H., Drowley, L., Barrett, I., Firth, M., Akerblad, P., Wolf, M. J., Bekiranov, S., Brautigan, D. L., Wang, Q.-D., & Saucerman, J. J. (2019). High-content phenotypic assay for proliferation of human iPSC-derived cardiomyocytes identifies L-type calcium channels as targets. Journal of Molecular and Cellular Cardiology, 127, 204–214. https://doi.org/10.1016/j.yjmcc.2018.12.015

Xin, M., Kim, Y., Sutherland, L. B., Murakami, M., Qi, X., McAnally, J., Porrello, E. R., Mahmoud, A. I., Tan, W., Shelton, J. M., Richardson, J. A., Sadek, H. A., Bassel-Duby, R., & Olson, E. N. (2013). Hippo pathway effector Yap promotes cardiac regeneration. Proceedings of the National Academy of Sciences, 110(34), 13839–13844. https://doi.org/10.1073/pnas.1313192110

Xin, M., Kim, Y., Sutherland, L. B., Qi, X., McAnally, J., Schwartz, R. J., Richardson, J. A., Bassel-Duby, R., & Olson, E. N. (2011). Regulation of Insulin-Like Growth Factor Signaling by Yap Governs Cardiomyocyte Proliferation and Embryonic Heart Size. Science Signaling, 4(196). https://doi.org/10.1126/scisignal.2002278

Yester, J. W., & Kühn, B. (2017). Mechanisms of Cardiomyocyte Proliferation and Differentiation in Development and Regeneration. Current Cardiology Reports, 19(13). https://doi.org/10.1007/s11886-017-0826-1

Yutzey, K. E. (2017). Cardiomyocyte Proliferation: Teaching an old dogma new tricks. Circulation Research, 120(4), 627–629. https://doi.org/10.1161/CIRCRESAHA.116.310058

Zeigler, A. C., Richardson, W. J., Holmes, J. W., & Saucerman, J. J. (2016). A computational model of cardiac fibroblast signaling predicts context-dependent drivers of myofibroblast differentiation. Journal of Molecular and Cellular Cardiology, 94, 72–81. https://doi.org/10.1016/j.yjmcc.2016.03.008

Zhang, Y., Liu, T., Meyer, C. A., Eeckhoute, J., Johnson, D. S., Bernstein, B. E., Nusbaum, C., Myers, R. M., Brown, M., Li, W., & Liu, X. S. (2008). Model-based analysis of ChIP-Seq (MACS). Genome Biology, 9(9), R137. https://doi.org/10.1186/gb-2008-9-9-r137

Zhao, M.-T., Ye, S., Su, J., & Garg, V. (2020). Cardiomyocyte Proliferation and Maturation: Two Sides of the Same Coin for Heart Regeneration. Frontiers in Cell and Developmental Biology, 8, 594226. https://doi.org/10.3389/fcell.2020.594226

Zheng, M., Jacob, J., Hung, S.-H., & Wang, J. (2020). The Hippo Pathway in Cardiac Regeneration and Homeostasis: New Perspectives for Cell-Free Therapy in the Injured Heart. Biomolecules, 10(7), E1024. https://doi.org/10.3390/biom10071024

