## Supplemental Material for "Dynamic map illuminates Hippo to cMyc module crosstalk driving cardiomyocyte proliferation"

Figure S1: Simulated activation of the cardiomyocyte proliferation network

Figure S2: Network-wide sensitivity analysis

Figure S3: Diminished sensitivity matrix with top influenced and influential nodes

Figure S4: Network robustness to variations in model parameters

Figure S5: Comparison of topological features versus influence

Figure S6: Virtual knockdown screen for regulators of cytokinesis

**Figure S1: Simulated activation of the cardiomyocyte proliferation network.**

The steady-state response of the network at baseline conditions.

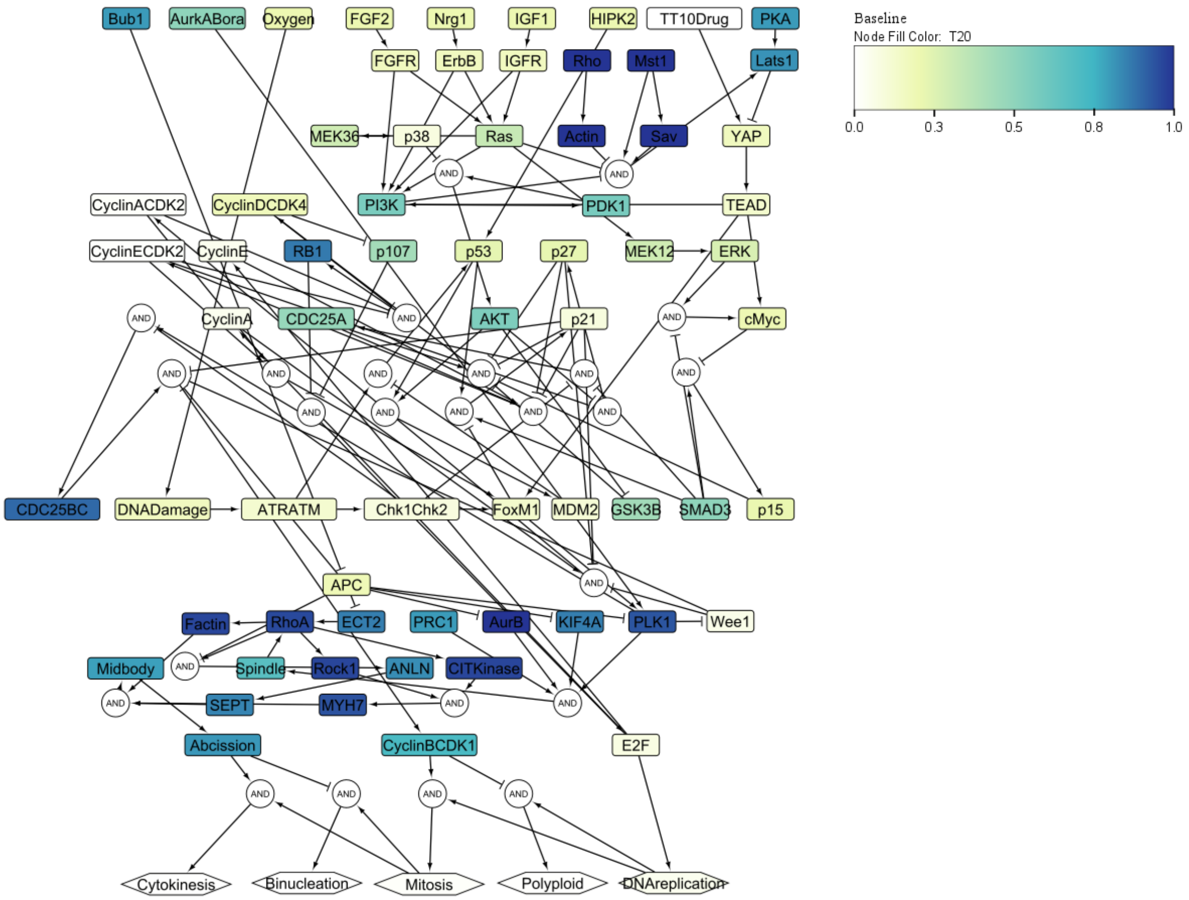

**Figure S2: Network-wide sensitivity analysis.**

Sensitivity analysis showing the effect of knockdown of each node on all other nodes in the model, in the baseline context. The columns of the matrix shown represent a simulation in which a node was knocked down and the change in activation of every other node in the network was measured. The top 30 most influential nodes (columns) and top 30 most sensitive nodes (rows) for the high Nrg1 context is shown in Fig. 3A. Blue indicates a decrease in activity with the knockdown of a node and red indicates an increase in activity with knockdown of the node.

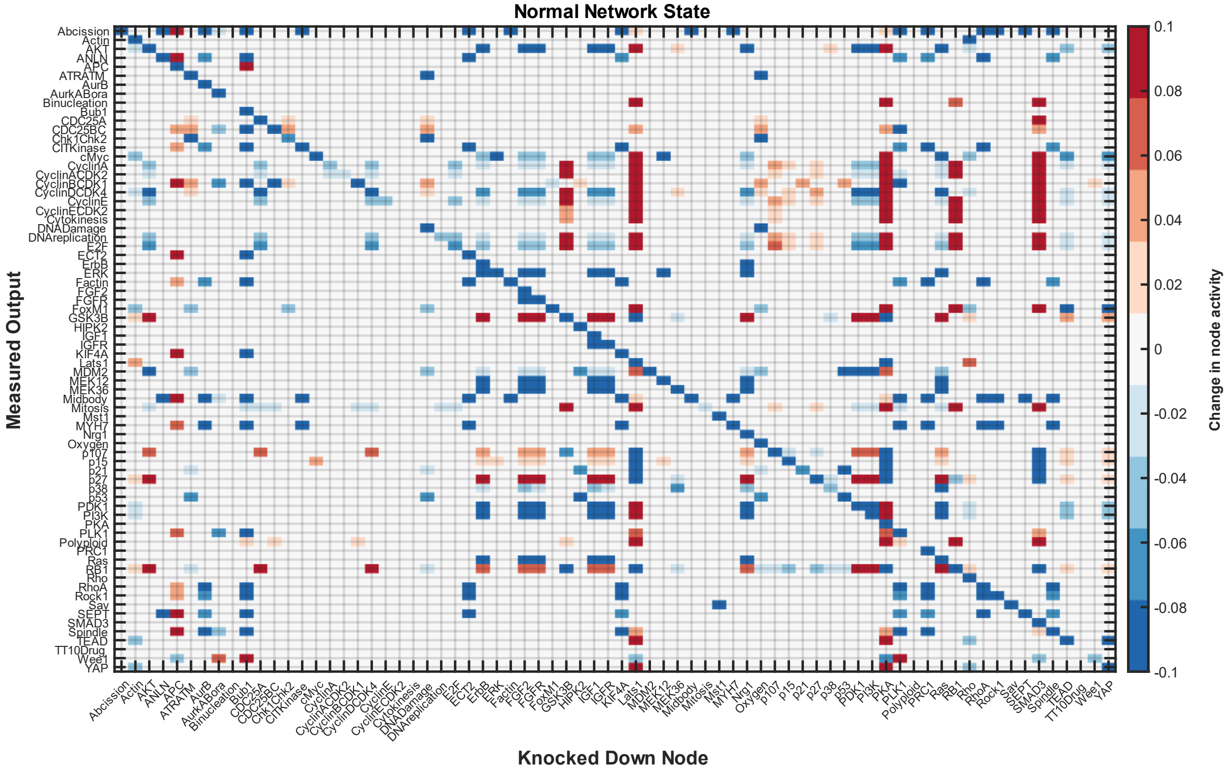

**Figure S3: Diminished sensitivity matrix with top influenced and influential nodes.** The top 25 most influential nodes (columns) and top 25 most sensitive nodes (rows), for the baseline (A), increased Nrg1 (B), and increased YAP (C).

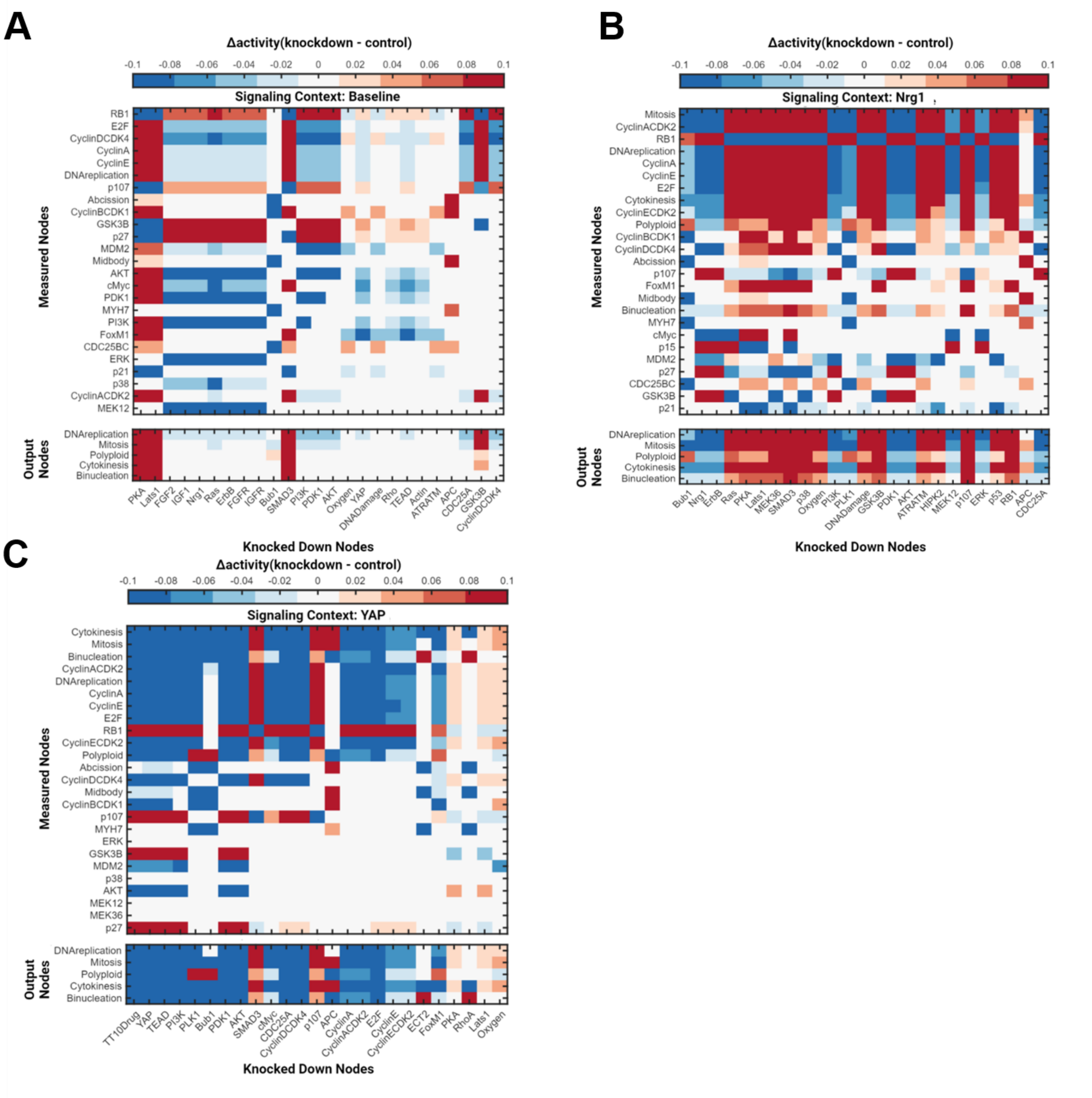

**Figure S4: Network robustness to variations in model parameters.**

For each parameter tested, parameters ($Y_{max}$ , w, and EC50) were varied within a uniform distribution of width 5%-50% of the original value. This uniform distribution was sampled 100 times for each parameter, and the simulations were run to compare model predictions with literature observations, using a validation threshold of 5% absolute change.

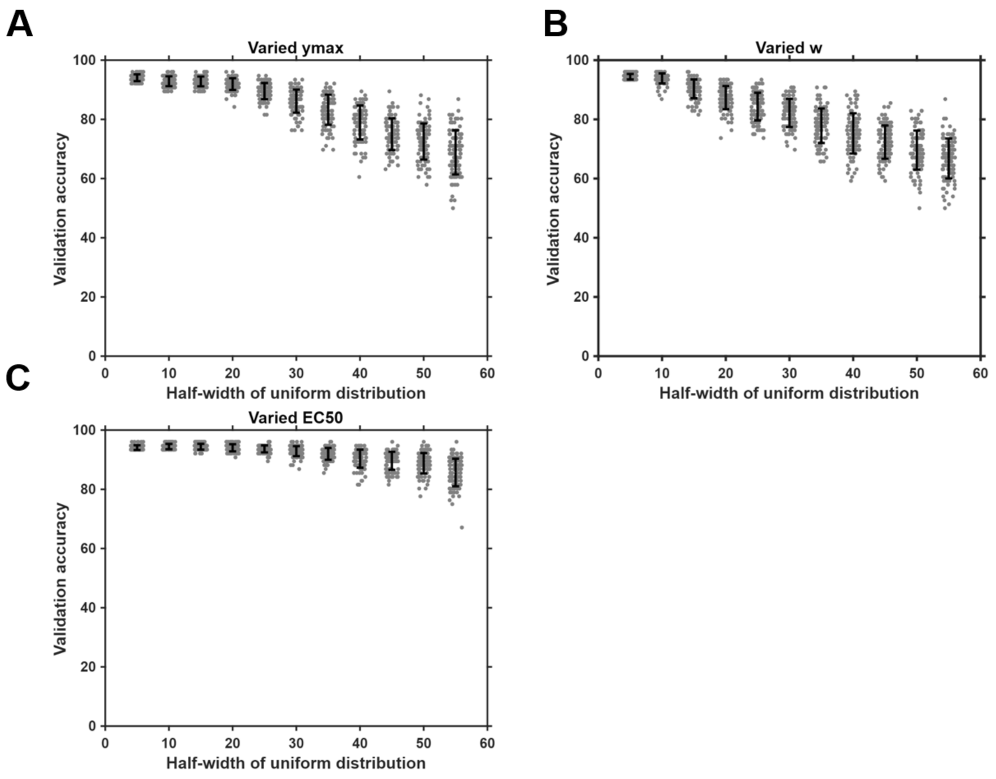

**Figure S5: Comparison of topological features versus influence.**

Eight topological features that summarize network structure (**Table S1**) were correlated against the influence and sensitivity metrics from sensitivity analysis under basal (A), high Nrg1 (B), and high YAP (C) conditions. Eccentricity and average shortest path were the most significant predictors of both influence and sensitivity in the high Nrg1 context. This differed from previous modeling of the fibroblast network, which showed betweenness centrality and degree as the most significant predictors of influence and sensitivity.

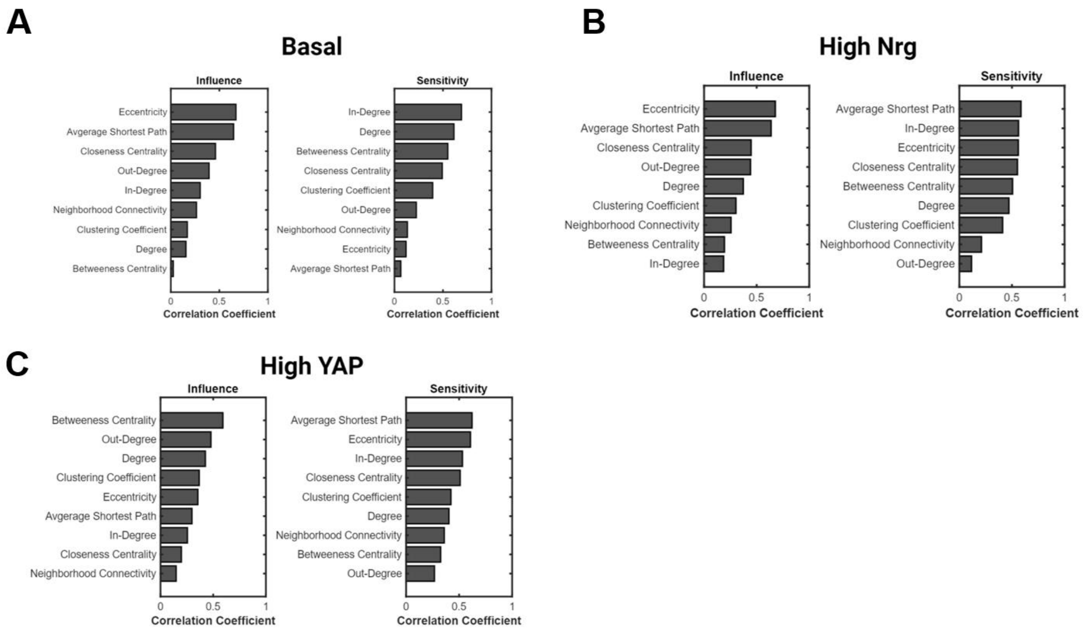

**Tables S1: Definitions of network topology metrics**

Each topological feature as defined by Assenov et al in the NetworkAnalyzer plugin for Cytoscape used for topological analysis of the network [16].

| Topology Metric | Defintition |
|---|---|
| betweenness centrality | number of shortest paths from all nodes to all others that pass through node n |
| out-degree | number of edges that exit node n |
| in-degree | number of edges that enter node n |
| eccentricity | maximal length of a shortest path between node n and any other node in the network |
| neighborhood connectivity | average number of neighbors of all neighbors of node n |
| average shortest path length | average shortest path between node n and any other node n |
| closeness centrality | the reciporical of the average shortest path length from node n (interpreted as a measure of how quickly information spreads from node n) |
| clustering coefficient | measure of the degree to which node n's neighbors form a complete graph |

**Figure S6: Virtual knockdown screen for regulators of cytokinesis**

A total knockout screen for cytokinesis in the basal or normal state (A), high Nrg1 (B), and high YAP (C). This analysis shows how sensitive cytokinesis is to the knockdown of all nodes of the model.

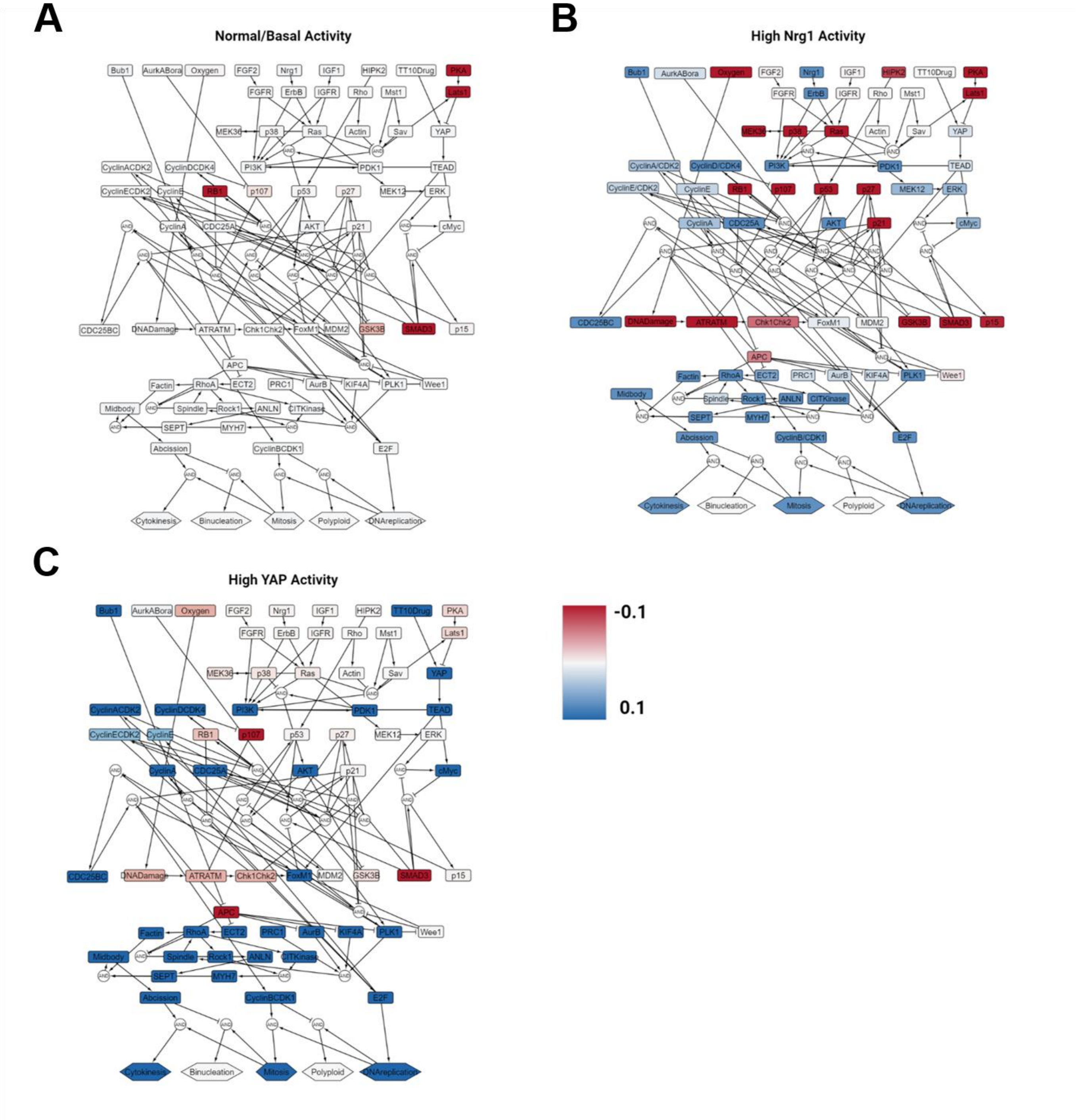
